## Supplementary information.pdf for "Assessing nanobody interaction with SARS-CoV-2 Nsp9"

### Supporting Information

**S1 File. Normalized attenuation ($A_N$) and relative error definitions.**

$A_N$ values were calculated according to:

$$A_N^k = \left(2 - \frac{\iota_p^k}{\iota_d^k}\right)$$

where the running index $k$ refers to the $k$th residue amide signal and the $\iota_{p,d}^k$ values are the corresponding auto-scaled intensities of each signal in the presence (subscript $p$ for paramagnetic) and absence (subscript $d$ for diamagnetic) of paramagnetic probe, defined as:

$$\iota_{p,d}^k = \frac{I_{p,d}^k}{\frac{1}{n}\sum_{k=1}^{n} I_{p,d}^k}$$

with $n$ being the total number of measured signals and $I$ denoting the signal intensities. From the latter equation, it is seen that the scaling factor is simply the mean value over the $n$ residues whose signal intensity can be estimated. The mean value of the individual auto-scaled intensities ($\iota_{p,d}^k$) is thus unitary, by definition [18-20]. Therefore, values of $A_N$ above or below unity identify larger or smaller attenuations, respectively, with respect to the normalized average attenuation.

From the previous definitions, the error on the individual $A_N$ values can be calculated as:

$$\Delta A_N^k = A_N^k \times \sqrt{\left[\frac{\Delta I_p^k}{I_p^k}\right]^2 + \left[\frac{\Delta I_d^k}{I_d^k}\right]^2 + \left[\frac{\frac{1}{n^2}\sum\left(\Delta I_p^k\right)^2}{\left(I_p^{av}\right)^2}\right] + \left[\frac{\frac{1}{n^2}\sum\left(\Delta I_d^k\right)^2}{\left(I_d^{av}\right)^2}\right]}$$

where the first two terms under the square root represent the error on the relative intensity value of the $k$th residue signal, that is, the signal intensity ratio in the presence and absence of nitroxide, the $\Delta I$ are the experimental intensity uncertainties obtained from the individual peak signal-to-noise value, and the superscript *av* stands for average.

**S1 Table. Peak loss and attenuation in $^{15}N$-$^{1}H$ HSQC spectra of SARS CoV-2 Nsp9 titrations with nanobodies.**

| | Peak loss | |
|---|---|---|
| **Nanobody : Nsp9 ratio** | **2NSP23[(a)]** | **2NSP90[(a)]** |
| 0.17:1 | - | - |
| 0.32:1 | - | *β4 (W53) l3-4 (50)* |
| 0.43:1 | *β1 (17) β3 (40, 46) β4 (52) β5 (67) β7 (89) l5-6 (69)* | *β2 (28) l5-6 (69)* |
| 0.54:1 | *β1 (12, 14) β2 (30, 31, 32, 33, 35) β4 (53, 54, 56) β5 (64, 66) β7 (87, 88) α1 (108) l1-2 (18, 19, 23) l3-4 (50) l7-α1 (91, 93)* | *β1 (12,17) β2 (30, 31, 32, 35) β3 (40, 41, 44, 45) β4 (52, 54, 56) β5 (67, 68) β7 (88, 89) α1 (108) l1-2 (18, 23) l7-α1 (93)* |
| 0.63:1 | *β1 (11, 13, 15) β2 (27, 29, 34) β3 ( 44, 45) β5 (65, 68) β6 (73) β7 (86, 90), α1 (95) l1-2 (22), l2-3 (37) l4-5 (58, 60, 62), l5-6 (70)* | *β1 (11, 13, 15) β2 (27, 29, 33, 33sc 34) β3 ( 38, 46) β4 (53sc) β5 (64, 66) β6 (74) α1 (95) l1-2 (19, 21, 22, 24) l2-3 (37) l4-5 (62, 63) l5-6 (70) l7-α1 (91)* |
| TOTAL | *β1 (**6**/**8**), β2 (**8**/**10**), β3 (**4**/**9**), β4 (**4**/**6**), β5 (**5**/**5**), β6 (**1**/**7**), β7 (**5**/**9**), α1 (**2**/**15**) l1-2 (**4**/**8**), l2-3 (**1**/**2**), l3-4 (**1**/**5**), l4-5 (**3**/**6**), l5-6 (**2**/**4**), l7-α1 (**2**/**4**)* | *β1 (**5**/**8**), β2 (**9**/**10**), β3 (**6**/**9**) β4 (**4**/**6**), β5 (**4**/**5**), β6 (**1**/**7**), β7 (**2**/**9**), α1 (**2**/**15**) l1-2 (**6**/**8**), l2-3 (**1**/**2**), l3-4 (**1**/**5**), l4-5 (**2**/**6**), l5-6 (**2**/**4**), l7-α1 (**2**/**4**)* |

| Fastest peak attenuation [(b)] | |
|---|---|
| **2NSP23** | **2NSP90** |
| *14, 17, 18, 19, 30, 32, 40, 46, 52, 54, 56, 67, 69, 89, 93, 108* | *13, 30, 31, 33, 41, 43, 51, 53, 54, 56, 67, 69, 108, 111* |

(a) The progressive loss of Nsp9 backbone or side-chain NH signals upon nanobody titration is reported with the indication, in parenthesis, of the residue number (italics) and the secondary structure location (β, α and *l* are strand, helix and intervening loop). The TOTAL rows report the counts of residue signals that are lost out of total number of residues in each secondary structure elements (numbers are in bold to avoid confusion with residue sequence numbers). The relative titrations were carried out at 278 K (2NSP23) and 276 K (2NSP90) to slow down the peak losses that were much more massive and fast at ambient temperature.

Although quite similar, the relative interactions of the two nanobodies exhibit slight differences for the involved epitopes, with 2NSP23 addressing a more extended surface on β7 strand, balanced in 2NSP90 interaction by small extensions of 1-2 loop and β3 strand. These differences map to epitopes e2 and e4 that are adjacent on each Nsp9 monomer contacted by 2NSP23 or 2NSP90 (see Fig. 3 of main text). Nsp9 sequence is reported below with the location of the secondary structure elements along the sequence highlighted in turquoise (β-strand) and yellow (α-helix). In addition, the nanobody sequences are also reported with the indication of the complementarity determining regions, CDR1 (blue), CDR2 (green) and CDR3 (red).

(b) The residue numbers (italics) with fastest attenuation upon nanobody titration are listed. The peaks with the steepest decrease in intensity were identified when the slope of their relative intensity attenuations was larger than the average attenuation slope increased by one standard deviation.

```
         10         20         30         40         50         60
NNELSPVALR QMSCAAGTTQ TACTDDNALA YYNTTKGGRF VLALLSDLQD LKWARFPKSD
             β1                     β2          β3                β4
         70         80         90        100        110
GTGTIYTELE PPCRFVTDTP KGPKVKYLYF IKGLNNLNRG MVLGSLAATV RLQ
      β5        β6          β7                 α1
```

#### 2NSP23:

```
         10         20         30         40         50         60
QVQLQESGGG LVQPGGSLRL SCAASGLAFS MYTMGWFRQA PGKEREFVAM IISSGDSTDY
         70         80         90        100        110        120
ADSVKGRFTI SRDNGKNTVY LQMDSLKPED TAVYYCAAPK FRYYFSTSPG DFDSWGQGTQ
        130        140
VTVSSAAAYP YDVPDYGSHH HHHH
```

#### 2NSP90:

```
         10         20         30         40         50         60
QVQLQESGGG LVQTGDSLRL SCAVSGRTFS TYSVGWFRQA PGKEREFVAL RWSGGTTYYA
         70         80         90        100        110        120
DSVVGRFTVS RDNAKNTVYL EMNSLKPEDT AVYYCAADRG SGSYSPTYRW DYWGQGTQVT
        130        140
VSSAAAYPYD VPDYGSHHHH HH
```

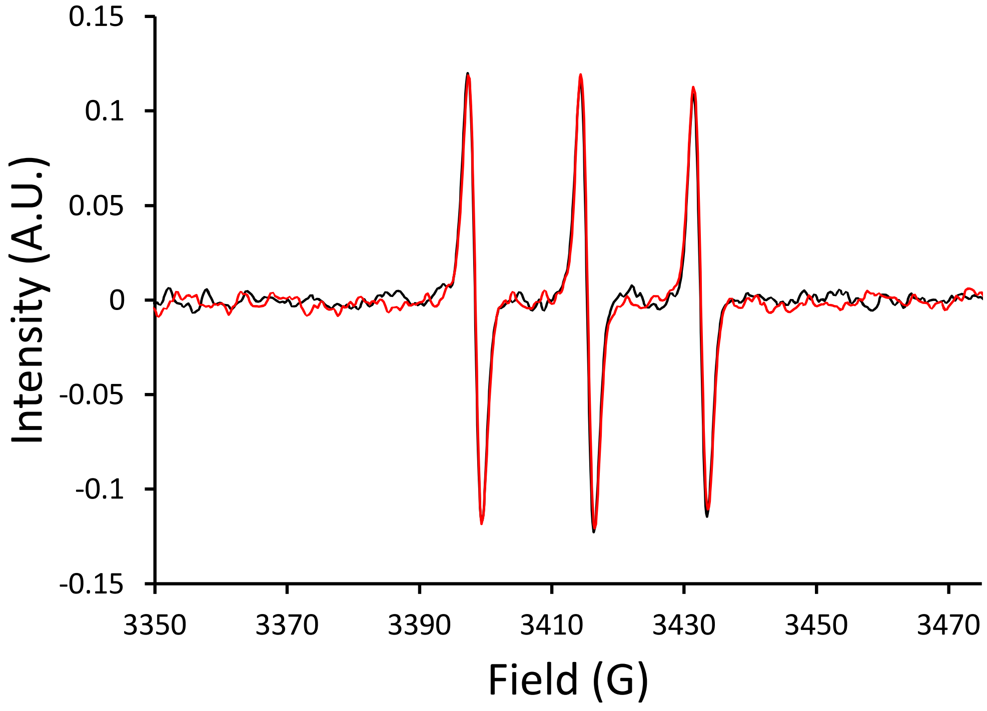


**S1 Figure. ESR spectra overlay.** The ESR spectra of 20 µM TEMPOL alone (black trace) and with 20µM SARS-CoV-2 Nsp9 + 20µM 2NSP90 (red trace) superimpose very well, confirming the invariance of the nitroxide dynamic regime in the presence of the proteins and hence the absence of specific tight interactions, consistently with the $\tau_c$ values reported in the main text that were calculated from signal spacings and amplitudes [18-20].

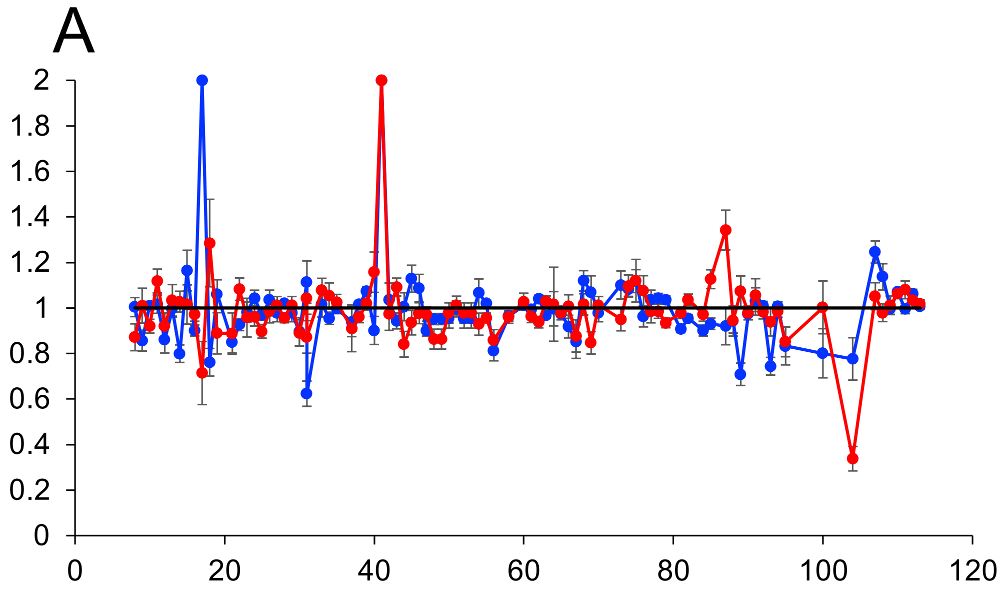


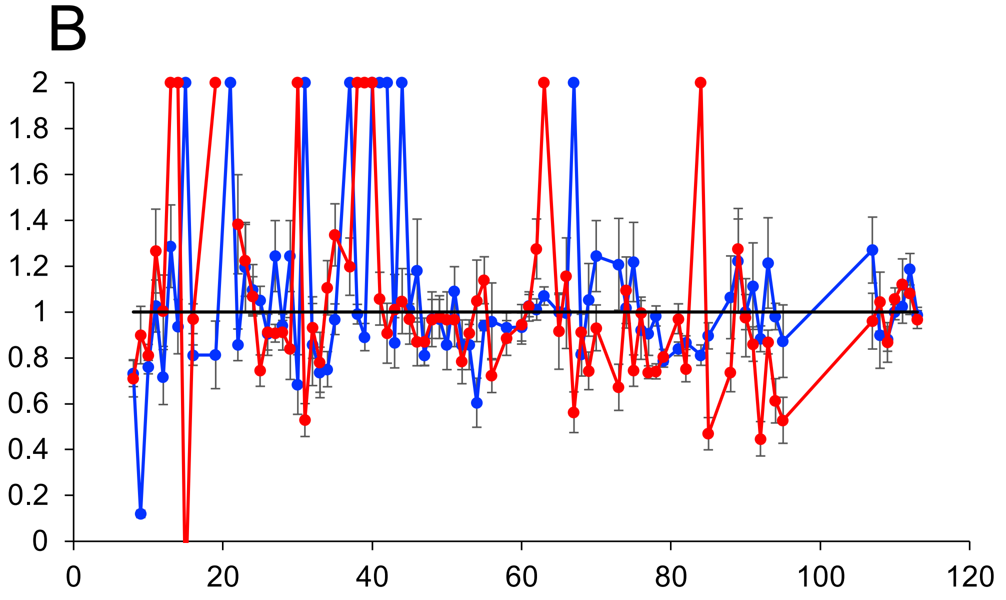


**S2 Figure. Normalized attenuation ($A_N$) pattern of Nsp9 – 2NSP90 system.** Overlay plot of the $A_N$ values obtained from $^1H$-$^{15}N$ HSQC spectra of 18 µM SARS-CoV-2 Nsp9 in the absence (A) and presence (B) of 5.6 µM 2NSP90 as determined by 1.0 mM Tempol at 298 K with a relaxation delay of 0.3 s (blue, nonequilibrium condition) and 3 s (red, equilibrium condition).

The data for segments The data for segments and residues 1−7, 86-87, 96−106, 20, 36, and 59 and the relative abscissa points are not reported because of the absence of the corresponding signals from the spectra. The locations of prolines (devoid of NH) are also skipped on the abscissa axis.

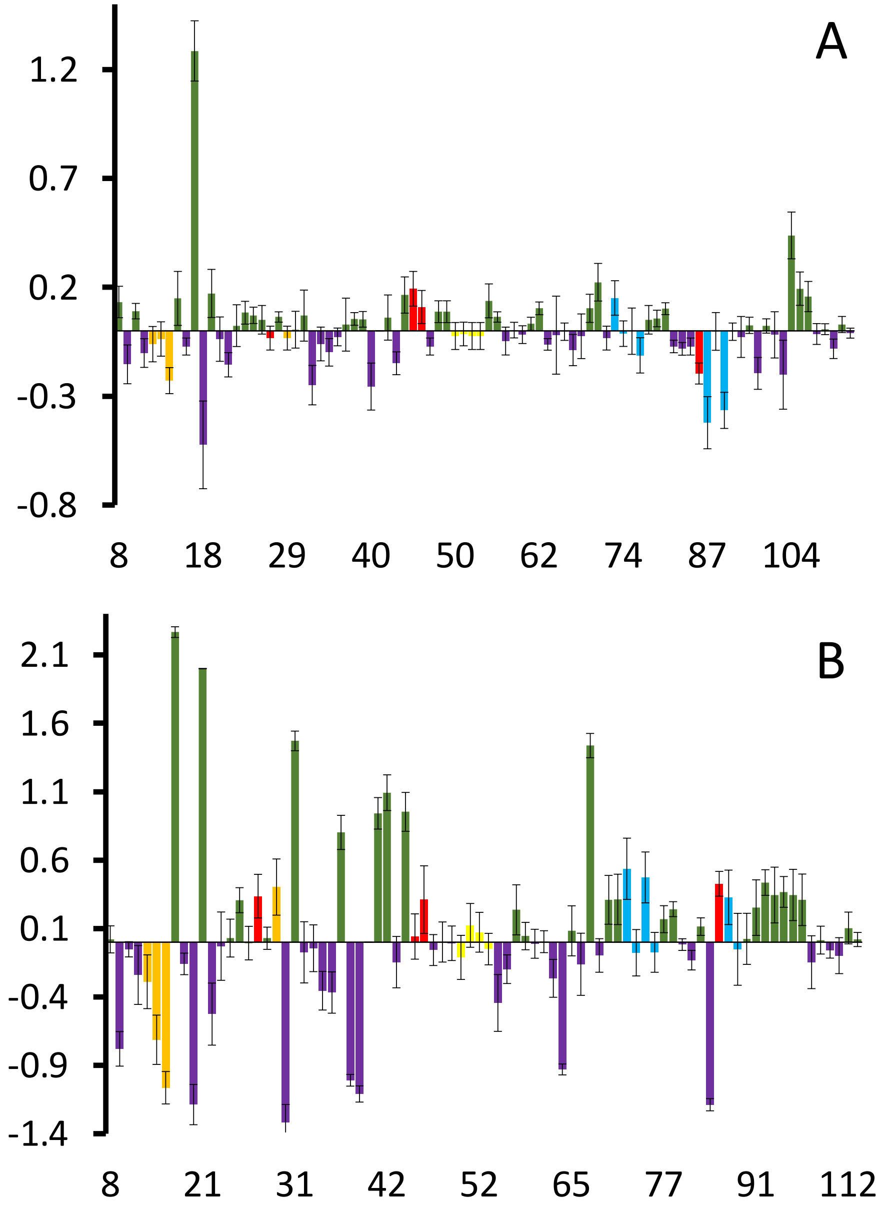


**S3 Figure. Nsp9 normalized attenuation differences.** Bar plot of {$A_N$[off] − $A_N$[eq]} differences highlighting the locations of the Type I pattern (green bars) and Type II pattern (purple bars) for 18 μM Nsp9 alone (A) and in the presence of 5.6 μM 2NSP90 (B). Same plot as Fig. 1 of main text without expansion truncation. Refer to Fig. 2 caption for color code and other information.

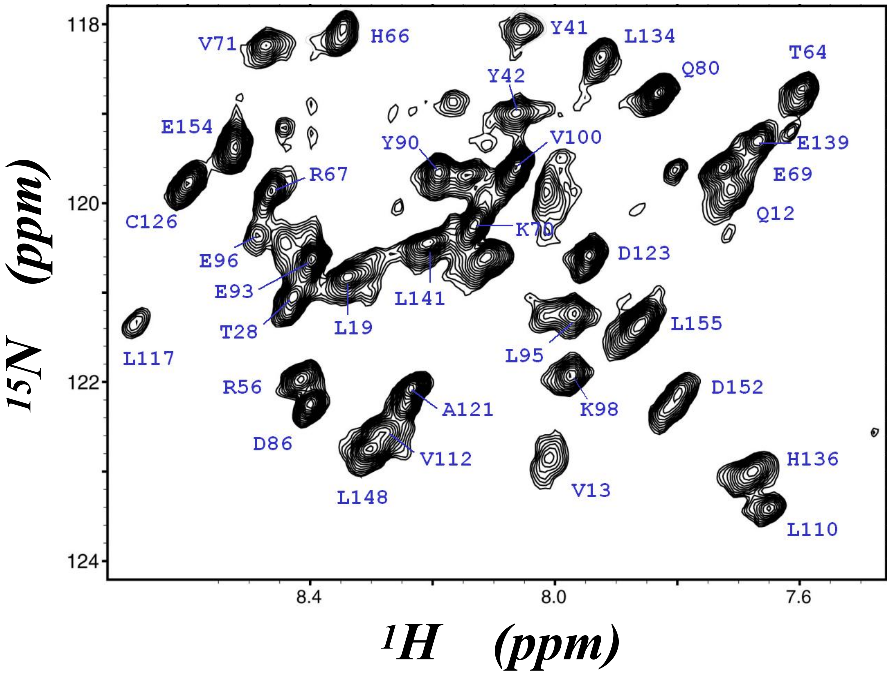


**S4 Figure. High molecular weight protein NMR.** $^{15}N$-$^{1}H$ HSQC spectrum U ($^{15}N$, $^{13}C$) 70% $^{2}H$ EMILIN1 C1q domain, a 52 kDa stable homotrimer [51]. The spectrum was obtained at 17.6 T (750 MHz $^{1}H$ frequency) and 310 K. The most crowded spectral region shown in the map benefits from 70% $^{2}H$ labeling, but a lower resolution spectrum is also observed with the fully protonated sample.
